# Scaling and merging macromolecular diffuse scattering with *mdx2*

**DOI:** 10.1101/2024.01.16.575887

**Authors:** Steve P. Meisburgera, Nozomi Andob

## Abstract

Diffuse scattering is a promising method to gain additional insight into protein dynamics from macro-molecular crystallography (MX) experiments. Bragg intensities yield the average electron density, while the diffuse scattering can be processed to obtain a three-dimensional reciprocal space map, that is further analyzed to determine correlated motion. To make diffuse scattering techniques more accessible, we have created software for data processing called *mdx2* that is both convenient to use and simple to extend and modify. *Mdx2* is written in Python, and it interfaces with *DIALS* to implement self-contained data reduction workflows. Data are stored in NeXusformat for software interchange and convenient visualization. *Mdx2* can be run on the command line or imported as a package, for instance to encapsulate a complete workflow in a Jupyter notebook for reproducible computing and education. Here, we describe *mdx2* version 1.0, a new release incorporating state-of-the-art techniques for data reduction. We describe the implementation of a complete multi-crystal scaling and merging workflow, and test the methods using a high-redundancy dataset from cubic insulin. We show that redundancy can be leveraged during scaling to correct systematic errors, and obtain accurate and reproducible measurements of weak diffuse signals.

**Synopsis:** *Mdx2* is a Python toolkit for processing diffuse scattering data from macromolecular crystals. We describe multi-crystal scaling and merging procedures implemented in the latest version of *mdx2*. A high-redundancy dataset from cubic insulin is processed to reveal weak scattering features.

## 1 Introduction

Diffuse scattering is the continuous pattern in the background of X-ray diffraction images from crystals^1^. It occurs whenever disorder is present, and significant motion is possible in macromolecular crystals; a typical protein crystal has 30-70% solvent, comparable to the crowded environment of cells^2^. Recently, the recognition that experimental data on protein dynamics are needed to understand their function has led to renewed interest in diffuse scattering^3^. In particular, diffuse scattering encodes unique information on the correlated displacements of pairs of atoms in the crystal^4^. Such correlated motions result, for instance, from the thermally-excited breathing motions of proteins, which have long been implicated in mechanisms of allostery and catalysis, but are challenging to study experimentally^5^.

A quantitative analysis of diffuse scattering requires carefully measured and processed data. In the reciprocal space mapping technique, diffraction data from one or more crystals in multiple orientations are combined to reconstruct the continuous, three-dimensional scattering pattern^6^. The state-of-the-art in reciprocal space mapping has evolved rapidly in recent years, driven by several technological advances. The availability of direct X-ray detectors at synchrotrons^7^ has enabled new data collection strategies to maximize data quality through fine *ϕ*-slicing^8^ and averaging many redundant observations^9^. The ideal properties of these detectors allow simultaneous measurement of Bragg and diffuse scattering in the same dataset^4,10^. New data processing tools, including *Lunus*^11^,*EVAL*^12^, *Diffuse*^13^, and *mdx-lib*^14^, have enabled reconstruction of the three-dimensional diffuse scattering patterns in reciprocal space. These patterns can be compared quantitatively with various models^15–17^.

By using these newly available tools, the field has reached a greater understanding of how different kinds of disorder contribute to diffuse patterns^3,13,14,18–20^. A key realization has been that atomic motions in protein crystals have correlations spanning a large range of distances, for instance from local atomic vibrations correlated over a few bond lengths, to wave-like excitations of the crystal lattice that may be correlated over many unit cells. As a consequence of the large range of length scales, diffuse patterns must be measured on a fine scale (i.e., oversampled with respect to the reciprocal lattice) in order to account for all of the observed atomic motion^14^. Moreover, in reciprocal space, the signals from the short- and long-ranged correlations are superimposed, and thus precise measurements are required in order to disentangle the signals from protein motions of interest^20^. Techniques to improve accuracy and precision have been developed, including experimental background measurement and subtraction^21^, procedures to correct for systematic errors during scaling^13,14^, and the development of new statistical indicators of data quality^22^ and model-data agreement^20^.

As the macromolecular diffuse scattering field grows beyond the community of methods developers, it is important to build data processing software that is easy to use, embodies best practices, promotes reproduciblility in research, and invites new communities of users and developers through documentation and tutorials. To address this community need, we developed *mdx2*, a Python package for processing macromolecular diffuse scattering data. A development version (0.3) has been available since 2022 with basic functionality suitable for education and preliminary data processing^6^. *Mdx2* is the successor to *mdx-lib*, a MATLAB library for diffuse scattering that has been used by the authors since 2016, and it re-implements many of its successful algorithms. Unlike *mdx-lib, mdx2* is designed to interface closely with the Bragg data processing program *DIALS*^23^. It features a user-friendly command-line interface, also inspired by *DIALS*, and stores intermediate data and metadata in standardized, self-describing NeXus-formatted hdf5 files^24^. The NeXus format was chosen to facilitate interchange with other software, such as the *NeXpy* graphical user interface^25^, or Jupyter notebooks via the *nexusformat* package. Although successful, the development version of *mdx2* lacked key features needed to handle large datasets produced in modern, high-redundancy, fine *ϕ*-sliced data collection, including support for multi-crystal scaling and parallel processing.

Here, we describe *mdx2* version 1.0, the first release intended for research, which includes an implementation of the full scaling model from *mdx-lib*^14^ and features parallelized data reduction tasks for multi-CPU architectures. This article is organized as follows. First, we provide an overview of data processing workflows combining *DIALS* and *mdx2* and describe the tasks performed by each command-line program. Next, we introduce the scaling model and refinement algorithm from *mdx-lib*, and describe its re-implementation in Python. Finally, we demonstrate new capabilities in *mdx2* by processing a large, multi-crystal dataset from cubic insulin collected at room temperature. We take advantage of the ∼70-fold redundancy of this dataset – unprecedented for diffuse scattering – to examine alternative statistical measures of data quality, and demonstrate the reproducibile detection of very faint diffuse signals at high resolution.

### 1.1 Overview of data processing in *mdx2*

*Mdx2* is a software package for processing diffraction data to reconstruct an accurate, three-dimensional map of reciprocal space. *Mdx2* breaks the reconstruction process into multiple steps that can be chained together to process data from one or more crystals. In developing the command-line interface for *mdx2*, we have focused on data collected using the traditional rotation method where the background scattering is measured at each rotation angle. This method, particularly when applied to large crystals, can robustly separate the scattering of the protein crystal from that of background sources, such as from air and mounting materials, and it has yielded high quality maps in previous diffuse scattering studies^14,20^.

#### 1.1.1 Diffraction geometry refinement and data import

The guiding philosophy of *mdx2* is that individual processing steps should consist of numerical algorithms that do not depend on experimental or crystallographic details. In practice, this means that the “import” steps in *mdx2* use crystallographic libraries (*dxtbx*^26^ and *cctbx*^27^) in order to pre-compute all necessary information for subsequent processing steps. For example, the symmetry operators for the space group are retrieved and converted to matrix form (Fig. 1A, step 6), and raw diffraction image data are re-compressed in a standard format (Fig. 1A, step 7). This approach has several advantages. First, standardization of data formats makes it easier to optimize the performance of algorithms, particularly when parallel processing is considered. Second, custom algorithms can be added easily, by developers or users, because they do not require specialized libraries or expertise in crystallographic computing. Finally, the pre-computed correction factors and detector data can be inspected directly before any processing occurs, which has significant value for education and exploratory data analysis^6^.

**Figure 1:**
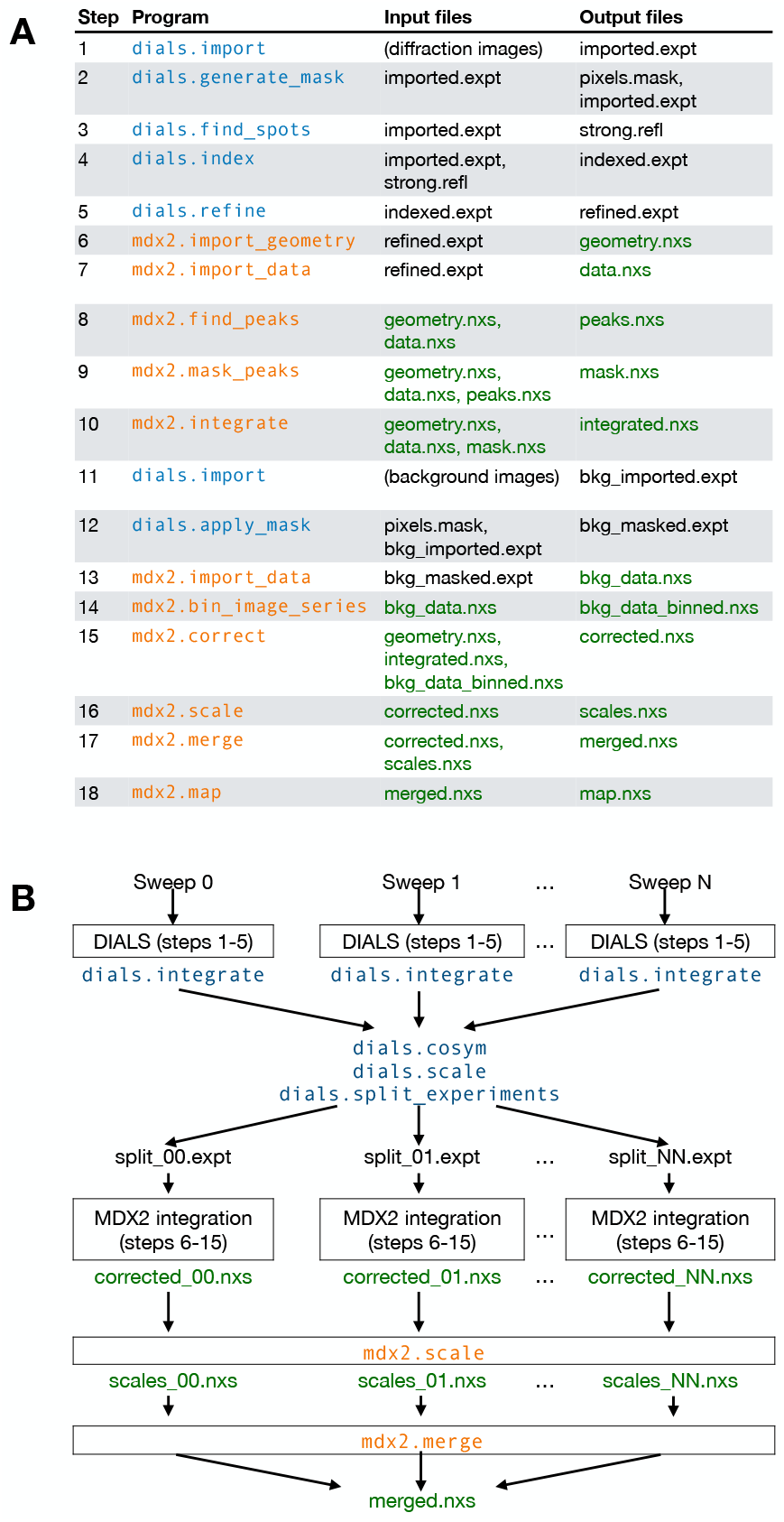
Data processing workflows. (**A**) Sequence of *DIALS* (blue) and *mdx2* (orange) command-line programs to process a single-sweep dataset with experimental background corrections. Diffraction geometry is refined by indexing Bragg peaks (steps 1-5) and the diffuse scattering is then integrated on a three-dimensional grid (steps 6-10). Background corrections are created (steps 11-14), and the integrated data are corrected, scaled, and merged (steps 13-18). NeXus-formatted hdf5 files (.nxs extension, green text) are used by *mdx2* for data and metadata exchange. See Methods and Meisburger & Ando [6] for details of each program. (**B**) Modified workflow for multi-sweep data. Each sweep is integrated independently in *DIALS*, then combined to resolve indexing ambiguities with *dials.cosym*^28^. Sweeps are processed independently in *mdx2* through the correct step, then the scaling model is refined globally, and sweeps are merged.

In a complete single-crystal workflow^6^, *DIALS* is first used to index the diffraction patterns and refine the detector geometry (Fig. 1A, steps 1-5). This step is critical in order to accurately assign each pixel to a location in three-dimensional reciprocal space (fractional Miller indices *h, k*, and *l*). A beamstop mask is also created (step 2, Fig. 1A), which will be carried over into diffuse processing. Although further processing in *DIALS* is not a necessary condition for proceeding to *mdx2*, integration and scaling should be performed to assess the overall Bragg data quality and to detect potential issues, such as radiation damage. Ultimately, the structure solved from the Bragg data will be combined with diffuse data to build a self-consistent disorder model.

Once Bragg data processing is completed, the geometry model from *DIALS* is imported into *mdx2* (step 6, Fig. 1A). All geometric corrections are pre-computed on a grid of points, including solid angle, air absorption, polarization, and detector efficiency. In addition, the fractional Miller indices are computed on a grid sampling detector position and rotation angle. Finally, symmetry information such as the Laue group operators, reciprocal space asymmetric unit, and reflection conditions are stored^6^. The pre-computed data and metadata are saved in NeXus format (geometry.nxs in Fig. 1A).

The data import step copies the detector image data from the raw format to NeXus format. Because *mdx2* and *DIALS* both use *dxtbx*^26^ to represent experimental geometry and read detector data, any image format compatible with *DIALS* can be read by *mdx2*. During import, the masks used in *DIALS* are applied to the image data (e.g., to remove bad pixels and the beamstop shadow). The data are stored as a three-dimensional array, or image stack, in NeXus format (data.nxs in Fig. 1A). Data are compressed in three-dimensional chunks so that small segments of the array can be read from disk without de-compressing the entire file, which is useful for certain algorithms and for parallel processing.

#### 1.1.2 Integration

In order to integrate the diffuse pattern, it is important to exclude pixels that are potentially contaminated by Bragg peaks. The strategy used in *mdx2* is to apply a mask wherever a Bragg peak is predicted to exist, regardless of its intensity. The mask has an ellipsoidal shape in reciprocal space (it is a function of the fractional coordinates Δ*hkl* relative to the nearest Bragg peak). Because the extent of a Bragg peak depends on many factors, including crystal mosaicity, beam divergence, and errors in the geometry model, the ellipsoidal shape is fit empirically in order to encompass the most intense features (step 8, Fig. 1A). First, the images are searched for all pixels with counts above a set threshold. For the dataset processed here, the threshold was set to 10 times the background scattering level of ∼2 photons per pixel. The detector location of each strong pixel is mapped to fractional Miller indices relative to the nearest whole integer. Then, an ellipsoidal Gaussian probability distribution is fit to the resulting point cloud. A binary mask is generated for each image excluding the Bragg peak regions (sept 9, Fig. 1A). The spatial extent of the Bragg peak region is set by a cutoff value expressed as a multiple of the Gaussian peak’s standard deviation. Any strong pixels (those exceeding a count threshold) not covered by the ellipsoidal masks are also flagged as outliers and masked out. Such outliers could arise, for instance, from broken detector pixels or diffraction from small salt crystals.

The integration task accumulates photon counts in a three-dimensional grid of voxels (Fig. 1A, step 10). The axes of the grid are aligned with the reciprocal lattice, and each voxel is assigned fractional Miller indices. When choosing a grid, voxel dimensions must evenly subdivide the reciprocal unit cell. In *mdx2*, any integer subdivision is allowed (its predecessor, *mdx-lib*, allowed only odd integer subdivisions). During integration, fractional Miller indices are calculated for each pixel according to the geometric model, and pixels are assigned to the nearest voxel. For each voxel, *mdx2* keeps track of the number of photon counts accumulated, the number of pixels contributing, and the mean position of the voxel in scan coordinates (the rotation angle and the location on the detector). The choice of grid spacing depends on the nature of the diffuse signal under investigation, as well as experimental considerations that potentially smear the diffuse pattern (see Meisburger & Ando [6]).

#### 1.1.3 Intensity corrections

After integration, the diffuse intensities are corrected for geometric effects and background scattering. Background corrections are prepared from a dataset taken with the crystal moved out of the beam^21^. These measurements include air scattering, scattering from mounting materials (such as capillaries), and shadows cast by the sample pin. The background images are first imported using *DIALS* and the beamstop mask is applied (Fig. 1A, steps 11-12). Then, the image stack is coarsened (binned down, potentially in terms of both pixels and frames) to reduce noise (step 14, Fig. 1A). Finally, the integrated photon counts from the crystal images are corrected using the background map and pre-computed geometric corrections. For each observation of a particular voxel, the correction factors are interpolated at the observation’s center position. The measured intensity and its uncertainty are estimated as follows:

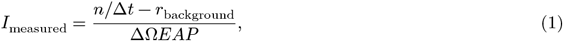

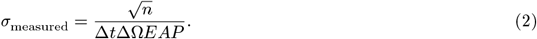

where *n* is the number of accumulated photons, Δ*t* is the cumulative exposure time per voxel (the number of pixels times the exposure time per image), *r*_background_ is the background scattering rate (photons per second), ΔΩ is the solid angle per pixel, *E* is the detector quantum efficiency, *A* is the transmission factor of air in the diffracted beam path, and *P* is the polarization factor^6^. The Poisson error due to background subtraction is reduced because of binning, and is neglected in Eq. 2.

#### 1.1.4 Scaling, merging, and visualization

Generally, a diffraction dataset contains redundant observations, either because the same region of reciprocal space is measured multiple times, or because of symmetries in the diffraction pattern. However, the equivalent observations may disagree as a result of systematic errors in the measurement, such as changes in the illuminated volume of the crystal as it is rotated. In *mdx2*, the parameters of a scaling model are refined in order to minimize the discrepancy between equivalent observations (Fig. 1A, step 16), thereby correcting the systematic errors. The model describes a correction applied to each observation that depends on its scan coordinates (such as rotation angle). The scaling model refinement procedures implemented in *mdx2* are detailed in Section 2.2. Multi-crystal workflows can also be orchestrated using *mdx2* (Fig. 1B). Scaling models for each rotation series are refined jointly.

Finally, the merging step outputs a table of fractional Miller indices, intensities, and estimated errors. In order to visualize the data in three dimensions, *mdx2* includes a mapping routine that symmetry-expands the data and places it in a three-dimensional array (Fig. 1A, step 18) suitable for plotting with *NeXpy*.

### 1.2 Scaling diffuse scattering data

The accuracy of diffuse scattering measurements may be limited by systematic errors. Artifacts commonly encountered during MX experiments include detector inhomogeneities, scattering from air in the beam path, scattering from the liquid or solid mounting materials that intercept the beam, and shadows cast by the sample and mounting materials. When an irregularly shaped crystal is partially illuminated by a small beam, which is often the case, the diffracting volume may also change during the scan, leading to a change in overall intensity. Additionally, some of the diffracted X-rays may be absorbed by the sample itself, leading to variations in intensity across the detector.

In Bragg data processing, *scaling* is the determination of scale factors for each independent observation that bring equivalent observations into agreement. The process of scaling involves fitting a model for the scale factors. Typically, the model parameters are physically motivated, and are parameterized to avoid over-fitting. Common effects included in scaling models include rotation-dependent changes in illuminated volume, absorption of diffracted X-rays, and B-factor decay due to radiation damage^29^.

Scaling diffuse scattering data has unique challenges. The diffuse signal of interest is typically a small variation (< 10%) on top of a largely homogeneous background^30^. Because of this, errors in scaling can very easily corrupt the small variations. In Bragg data, the local background is subtracted from each Bragg peak prior to integration, and thus changes in background scattering do not need to be corrected. In contrast, for diffuse scattering, the excess background scattering must be considered. Finally, since Bragg data and diffuse scattering come from the same crystal volume, the same scaling model should apply to both signals. In practice, it has not been possible to transfer the Bragg scaling model to diffuse scattering data, possibly because radiation damage-induced decay of Bragg intensitites is significant at ambient temperature. In the workflow presented here, Bragg and diffuse data are scaled independently.

In general, redundant observations are required in order to fit a scaling model. High redundancy (much greater than 2-fold) has significant advantages for diffuse scattering. Outliers can be more easily identified and elliminated, scaling model refinement is more robust, and systematic errors not included in the scaling model are more likely to be averaged out.

#### 1.2.1 Theoretical background

The scaling process minimizes the discrepancy between the intensity predicted by the scaling model and each corresponding observation. The discrepancy can be written as a *χ*^2^ statistic, which is minimized:

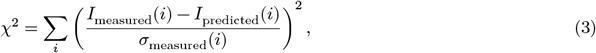

where the summation is over all observed voxels indexed by *i, I* is the intensity, and *σ* is the standard error of the measurement. The predicted intensity (*I*_predicted_) is initially unknown. For linear scaling models, the predicted intensity can be written schematically as a linear transformation of the “true” intensity *I*_0_ as follows:

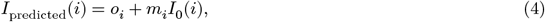

where *o*_i_ is the offset and *m*_i_ is the scale factor.

Combining Eq. 3 and Eq. 4, and rearranging terms,

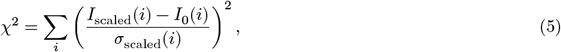

where 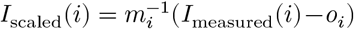 and 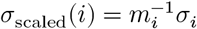 are the inverse-scaled observations and uncertainties.

The equation for optimally merging scaled intensities can be derived from Eq. 5 as the value of *I*_0_ that minimizes *χ*^2^, with the following result:

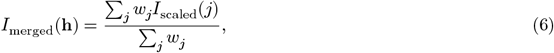

where *w*_*j*_ = [*σ*_scaled_(*j*)]^−2^ are the weights, and the sums run over all equivalent reflections *j* with Miller index **h** in the asymmetric unit of reciprocal space. The uncertainty is as follows:

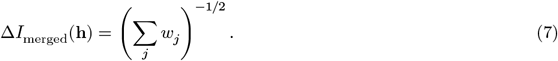

Both the scale model parameters and the merged intensity are unknown, and must be estimated simultaneously. This leads to a nonlinear optimization problem. An iterative method may be applied when the scale factors are linear functions of model parameters^31^. Repeated cycles of merging and scale model fitting converge toward the solution that minimizes *χ*^2^.

### 1.2.2 Scaling model refinement

The formalism for least-squares optimization can be expressed conveniently in matrix form^32^. Conventionally, *χ*^2^ is written as the L2-norm of a vector of residuals as follows:

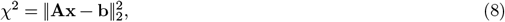

where **x** is a vector of unknowns, **b** is a known vector, and **A** is a matrix. The unknowns are the solution to the following linear equation:

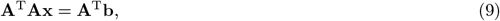

which can be solved directly by matrix inversion if the problem is not underdetermined. When the model itself is underdetermined by the data, or when there is a danger of overfitting, it is often useful to apply restraints to the model, such as smoothness. In the method of regularized least-squares, an additional term is added:

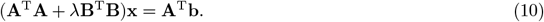

The matrix **B** is an operator that acts on the parameter vector **x** to calculate the quantity to minimize in an L2-norm sense. For instance, **B** may calculate a discrete representation of the second derivative in order to obtain solutions that are smooth^32^. The pre-factor *λ* is known as the regularization parameter, and it sets the trade-off between satisfying the restraints and goodness of fit.

As a concrete example, consider the general scaling model (Eq. 4) where the scale factor *m*_*i*_ depends only on the *ϕ* angle of the crystal. Let the model be parameterized by a set of scale factors (the parameters) at regular intervals, where the scales for each observation are calculated by linear interpolation between the control points. The linear interpolation can be expressed as a linear operator acting on a vector of parameters as follows:

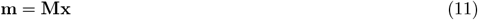

By algebraic manipulation of Eq. 5, it can be shown that the least-squares solution for **x** is obtained from Eq. 9 with the following substitutions:

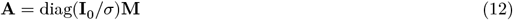

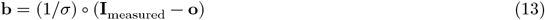

In the above notation, the *diag* function creates a diagonal matrix from its vector argument, and the open circle and forward slash (/) operators denote element-wise multiplication and division of vectors, respectively.

Based on this formalism, we have applied the following general procedure to derive more complex scaling models:

1. Write the complete scaling model in terms of its prediction for each observation given the “true” intensity.
2. Choose a convenient parameterization (typically linear intepolation on a grid of dimension 1-3).
3. For each scale factor and offset, derive a linear operator that calculates the value for each observation given the relevant coordinates of the observation and a vector of parameters.
4. Rearrange Eq. 5 to find the **A** matrix and **b** vector to be used in least-squares minimization.
5. Derive operators to use in regularization (typically enforcing a notion of smoothness)

## 2 Methods

### 2.1 Scaling model

We recently introduced a scaling model that accounts for common experimental artifacts in diffuse scattering data^14^. The model contains four physically-motivated parameters, labeled *a*-*d*, that describe a linear transformation of the true intensity *I*_0_ at each observation point *i* (Eq. 4), as follows:

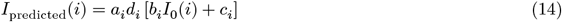

The three scale factors (*a, b*, and *d*) and offset term (*c*) are themselves functions of the coordinates for each observation: the rotation angle (*ϕ*), position on detector (*x, y*), and scattering vector magnitude (*s*). Each parameter’s coordinate dependence and its physical meaning are summarized in Table 1.

**Table 1:**
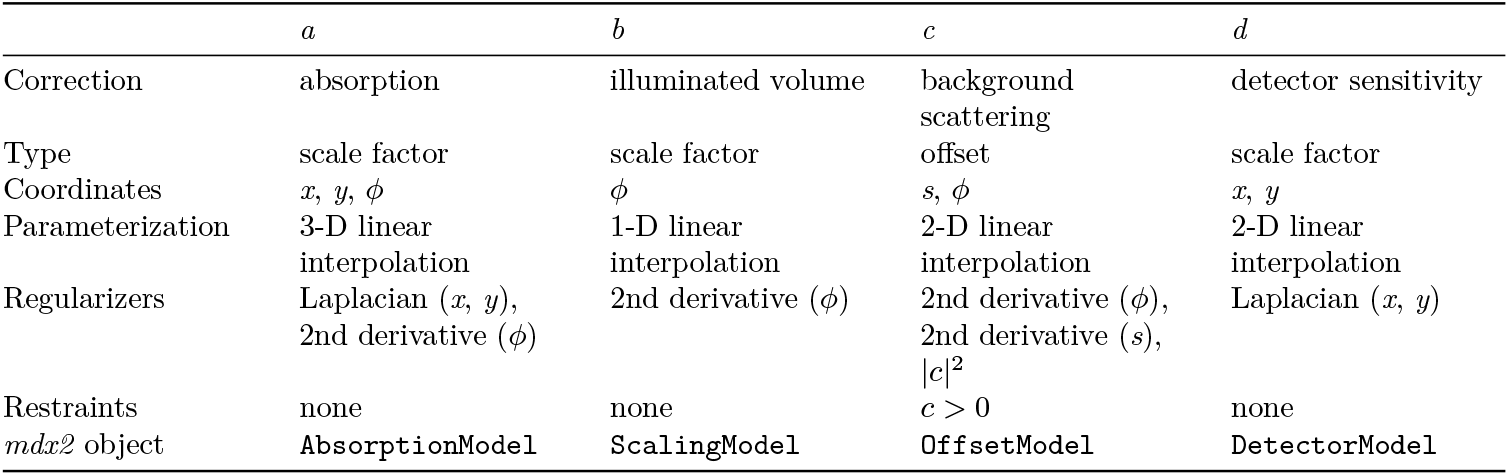
Scaling model parameters.

Numerically, the parameters are represented by discrete samples on a grid of dimension 1-3, and the value at each observation’s coordinates is computed by linear interpolation (Table 1). Interpolation is computed using a matrix operator^33^ acting on a vector of control points, as in Eq. 11. In one dimensional interpolation, the vector is simply an ordered list of the discrete samples. In higher dimensions, the grid of control points is vectorized, and the interpolation operators are constructed in a corresponding manner.

The general procedure to refine a given parameter is to alternate between two linear least-squares minimizations: first, to find the “true” intensity *I*_0_ that minimizes *χ*^2^ with the scaling model held constant (Eq. 6); and second, to find the scaling parameter values that minimize *χ*^2^ plus the regularizing terms (Eq. 10) given the previous value for the merged intensity. The procedure is repeated for all four parameters in the scaling model. To simplify the implementation, each least-squares minimization step is expressed in a common notation, where the **A** matrix and **b** vector appearing in Eq. 10 are derived for each parameter, and **M** represents the linear interpolation operator (Table 2).

**Table 2:** Least-squares problem for each scaling parameter.

| | $\mathbf{A}$ | $\mathbf{b}$ |
| --- | --- | --- |
| $a$ | $\text{diag}(\mathbf{d} \circ (\mathbf{b} \circ \mathbf{I}_0 + \mathbf{c})/\sigma)\mathbf{M}$ | $\mathbf{I}/\sigma$ |
| $b$ | $\text{diag}(\mathbf{d} \circ \mathbf{a} \circ \mathbf{I}_0/\sigma)\mathbf{M}$ | $(\mathbf{I} - \mathbf{a} \circ \mathbf{c} \circ \mathbf{d})/\sigma$ |
| $c$ | $\text{diag}(\mathbf{d} \circ \mathbf{a}/\sigma)\mathbf{M}$ | $(\mathbf{I} - \mathbf{a} \circ \mathbf{b} \circ \mathbf{d} \circ \mathbf{I}_0)/\sigma$ |
| $d$ | $\text{diag}(\mathbf{a} \circ (\mathbf{b} \circ \mathbf{I}_0 + \mathbf{c})/\sigma)\mathbf{M}$ | $\mathbf{I}/\sigma$ |

Each parameter has one or more associated regularizing operators to enforce a notion of smoothness (Table 1). We have chosen operators with straightforward **B**-matrix implementations^32^: a discrete 2nd derivative operator regularizes the *ϕ*- and *s*-dependence, and a two-dimensional discrete Laplacian operator regularizes the (*x, y*) dependent parameters. Each operator is weighted independently by a regularization parameter during least-squares minimization (*λ* in Eq. 10). Reasonable values for the regularization parameters are estimated by the fitting program (see Implementation, below), but in general they can be tuned by the user to control the smoothness of the solution based on an understanding of the experiment.

The offset parameter (*c*) has additional restraints and regularizers in order to make the correction as small as the data allow. This is important because an arbitrary offset may be added and subtracted from the merged intensity and the scaling model, respectively, leading to the same *χ*^2^ in Eq. 5. Physically, it is assumed that at least some of the data do not require offset corrections; i.e., the beam intersects only the crystal during at least some part of the scan, and the experimental background has been correctly subtracted. The offset correction is therefore always positive, and an additional regularization operator (an identity matrix) penalizes large values of the parameter. The positivity restraint is nonlinear and cannot be enforced using the regularized least-squares formalism of Eq. 10. Instead, the restraint is applied during iterative refinement of the offset parameter, an approach that is widely used for such restraints in the context of multivariate curve resolution^34,35^. After fitting the parameter values, any negative values are set to zero. If all of the points are positive, a constant is subtracted to make at least one of the values equal to zero. Then, the fit is repeated until reaching convergence (described further in Implementation, below).

### 2.2 Implementation of scaling model refinement in *mdx2*

The operations for scaling and merging described above were implemented in Python following the same modular approach that is used for other functions in *mdx2*. First, a library within *mdx2* called *scaling* was created that is essentially a toolkit from which algorithms may be built. The algorithm itself is implemented in the command-line function `mdx2.scale,` which imports the *scaling* library, handles file inputs and outputs, sets regularization parameters, and controls flow through the algorithm (such as order of refinement operations and halting conditions). The default behavior of `mdx2.scale` is to run a primitive scaling model where only the *b* term is refined. The full scaling model is activated using the flag `--mca2020,` which tells the program to mimic the behavior and default parameters from *mdx-lib*^14^. However, the algorithm can be customized at the command line using non-default parameters and individual terms in the scaling model can be disabled entirely if required. As in all *mdx2* command-line programs, the options and their default values are printed using the `--help` flag (Appendix B).

The *scaling* library contains a hierarchy of objects (Python classes) that build on one another to execute scaling operations for each type of parameter with minimal duplication of code. At the lowest level of the hierarchy are the linear interpolation objects (`InterpLin1, InterpLin2, and InterpLin3`) that compute sparse matrix representations of the linear interpolation operators in 1-3 dimensions, as well as the Laplacian and 2nd derivative operators used in regularization. Corresponding objects were previously implemented in the MATLAB library *mdx-lib*, and were translated directly into Python following the example in our *REGALS* package for small angle scattering data analysis^33^, which has both MATLAB and Python implementations of `InterpLin1.`

Each type of scaling parameter is stored using a corresponding Python object with a consistent interface (Table 1). The objects store parameter values on a grid of scan coordinates, and these values can be converted to/from NeXus data arrays for input and output.

The `ScaledData` object stores the unscaled observations, their reciprocal space indices, scan coordinates, and the current value of the overall scale and offset for each observation. It includes functions to facilitate selecting batches (i.e., single sweeps of data) via the Python iterator syntax, and for merging equivalent observations using Eq. 6.

Each scaling model object (Table 1) has a corresponding `ModelRefiner` object with a common interface. The function `calc_problem` returns the matrix-matrix and matrix-vector products used in least-squares minimization (Eq. 10 and Table 2), and the fit method performs the least-squares minimization step given the current merged intensity and scaling parameters. Regularized least squares is used with regularization parameters passed as arguments to the `fit` method. Each regularization parameter is re-normalized following the approach used in *REGALS*^33^, as follows:

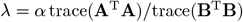

where *α* is the regularization parameter specified by the user, and *λ* is the pre-factor of the regularization term **B**^T^**B** (see Eq. 10). This re-normalization means that a certain value of *α* will produce similar results despite different grid spacings, dataset sizes, and noise levels in the data. In general, *α* ≪ 1 will favor minimization of *χ*^2^, while *α* ≫ 1 will favor minimization of the regularizer. For default values of *α*, see the Supporting Information.

### 2.3 Parallel processing in *mdx2*

Computationally-intensive processing steps in *mdx2* can be efficiently distributed over multiple processors if available. The image stack is divided among the workers in three-dimensional segments that are aligned with the hdf5 chunks specified in `mdx2.import_data.` In *mdx2* version 1.0, multiprocessing is available in `mdx2.import_data, mdx2.find_peaks,` mdx2.mask_peaks, and `mdx2.integrate.` This feature is activated by specifying the number of processors with the --nproc flag. Note that in `mdx2.import_data`, the performance is currently limited by the single-threaded hdf5 output file writer. In other cases, performance scales with the number of processors.

### 2.4 Experimental methods

Insulin crystals in the Zn-free cubic form were prepared following published protocols^36^. Insulin from bovine pancreas was purchased as a lyophilized powder (Sigma cat. no. I5500) and used without further purification. Drop volume, pH, and precipitant concentration were optimized to favor the growth of large single crystals (Table 3).

**Table 3:** Crystallization.

|  |  |
| --- | --- |
| Method | Sitting-drop vapor diffusion |
| Plate type | Cryschem 24 well (Hampton) |
| Temperature | 295 K |
| Protein concentration | 20 mg mL<sup>-1</sup> |
| Buffer composition of protein solution | 20 mM Na phosphate dibasic |
| Composition of reservoir solution | 300 mM Na phosphate pH 10.0, 10 mM Na EDTA |
| Volume and ratio of drop | 16 μL, 1:1 |
| Volume of reservoir | 500 mL |

X-ray diffraction data were collected at ambient temperature at the Cornell High Energy Synchrotron Source (CHESS) beamline F1 following diffuse scattering protocols introduced previously^14,21^. The beamline and data collection parameters are summarized in Table 4. At the synchrotron, two large insulin crystals were harvested from the the same drop of a crystallization tray using Kapton loops. The loops were immediately placed in plastic capillaries pre-loaded with reservoir solution to maintain hydration. In total, 17 datasets were collected from distinct locations on the two crystals using fine *ϕ*-slicing and low dose per frame. A total exposure time of 50 s for each location was chosen to limit radiation damage to tolerable levels, as judged by the B-factor decay obtained during scaling. For each crystal, a set of background scattering measurements were made by translating the crystal out of the beam along the rotation axis and collecting 360° of data at 1° s^-1^ while acquiring images at 1 Hz (1 s and 1° per frame).

**Table 4:** Data collection.

|  |  |
| --- | --- |
| X-ray source | Cornell High Energy Synchrotron Source (CHESS) beamline F1 |
| Wavelength | 0.9768 Å |
| Energy bandwidth ($\Delta E/E$) | ~0.05% |
| Beam size, profile | 100 μm diameter round, flat-top |
| Detector | Pilatus 3 6M, 1.0 mm Si sensor |
| Rotation rate | 1.0 deg. s<sup>-1</sup> |
| Rotation per image | 0.1 deg. |
| Temperature | 295 K |
| Crystal mount | 800 μm diameter loop (MiTeGen DT) inside PET capillary (MiTeGen Micro-RT) with 10 μL reservoir solution |
| Number of crystals | 2 |
| Number of diffraction images | 8500 |

## 3 Results and discussion

### 3.1 A high-redundancy room temperature dataset for diffuse scattering

To validate and test new capabilities of *mdx2*, we chose to process a room temperature multi-crystal dataset from cubic insulin. Diffraction data were collected from two nominally identical crystals (Table 4) using a multi-sweep strategy to distribute the dose (Fig. 2). To determine whether all frames collected are of consistent quality, we processed the Bragg data using a DIALS multi-crystal workflow (Fig. 1B). According to the B-factor decay model fit during scaling, the onset of radiation damage was immediate and it progressed in a similar manner at each location (Fig. 2, bottom axes). The change in overall B-factor during exposure ranged from 2-4 Å_2_ per 50 degrees of rotation, which is comparable to previously-reported diffuse scattering datasets from lysozyme^14,20^. The data processed to a resolution of 1.20 Å (Table 5). With each crystal processed separately, statistics such as *R*_pim_, mean *I / σI*, and *CC*_1/2_ indicate that the data are of excellent quality (Crystal 1 and Crystal 2, Table 5). These statistics improve further when the two crystal datasets are processed together (Crystals 1 & 2, Table 5), indicating that the crystals are highly isomorphous and of comparable quality. The final merged dataset has a multiplicity of 68.5, which far exceeds that of any macromolecular diffuse scattering dataset reported to date.

**Table 5:** Bragg data collection statistics. ^*^Values in parentheses refer to the highest resolution shell

|  | Crystals 1 & 2 | Crystal 1 | Crystal 2 |
| --- | --- | --- | --- |
| Crystal parameters |  |  |  |
| Space group | I 2<sub>1</sub> 3 |  |  |
| Cell dimensions (Å) | $a=b=c=79.519$ | | |
| Dataset statistics |  |  |  |
| Resolution (Å) | 28.11–1.20 (1.22–1.20)* |  |  |
| $CC_{1/2}$ | 1.000 (0.276) | 1.000 (0.115) | 1.000 (0.226) |
| $R_{\text{merge}}$ | 0.054 (2.236) | 0.058 (2.539) | 0.046 (1.758) |
| $R_{\text{pim}}$ | 0.006 (0.769) | 0.009 (1.113) | 0.007 (0.868) |
| Mean $I / \sigma I$ | 44.8 (0.6) | 28.9 (0.3) | 37.2 (0.6) |
| Completeness (%) | 99.81 (96.32) | 98.38 (77.68) | 99.08 (83.97) |
| Multiplicity | 68.5 (8.7) | 36.6 (5.7) | 32.5 (4.8) |
| Observations | 1795925 (10983) | 944563 (5742) | 846333 (5241) |
| Unique reflections | 26218 (1256) | 25842 (1013) | 26026 (1095) |

**Figure 2:**
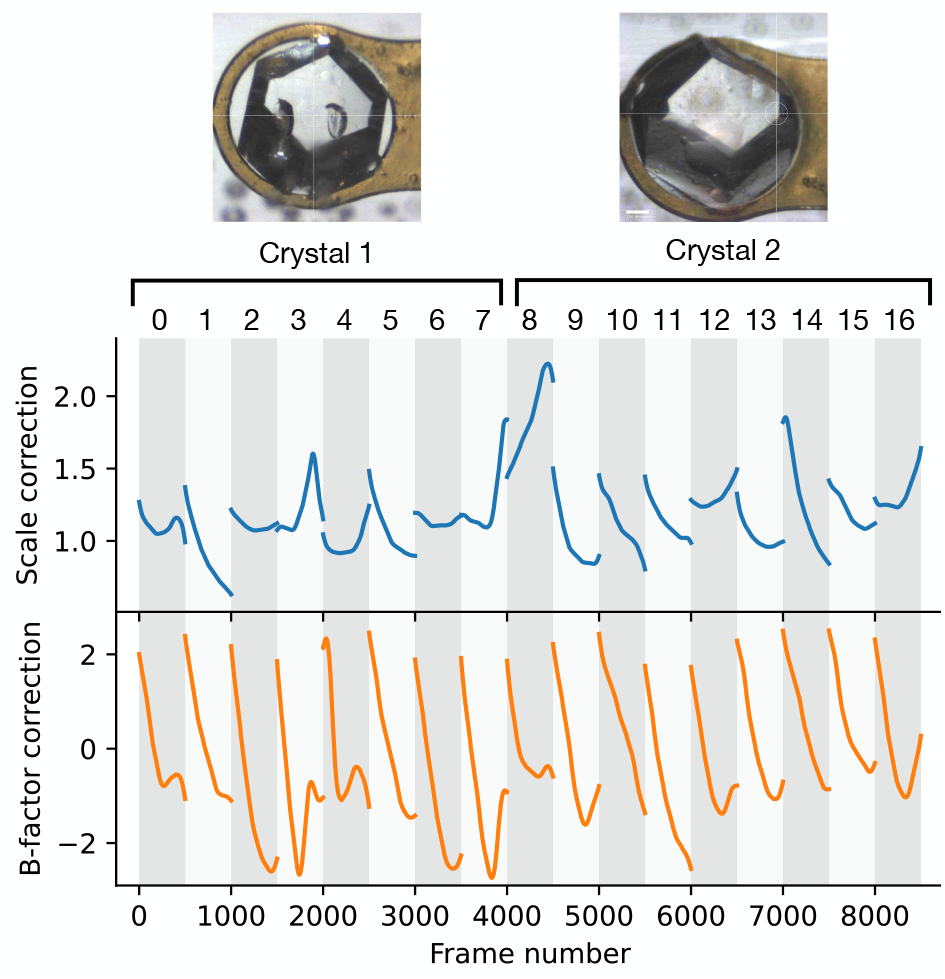
Bragg intensity scaling results from a multi-crystal insulin dataset. Rotation datasets of 50 degrees (500 frames each) were collected from 17 locations on two large insulin crystals (pictured). Each 50-degree wedge (sweep) spans the width of the white/gray vertical bars. Correction factors from global scaling in *DIALS* are shown for each wedge. The illuminated volume changes during rotation, as reflected in the overall scale factor (top axes, blue curves). B-factors show a characteristic linear decay with exposure time due to radiation damage (bottom axes, orange curves). The overall extent of radiation damage is consistent among the datasets.

### 3.2 Data reduction in *mdx2*

After initial processing in DIALS, each rotation dataset (sweep) was imported into *mdx2* (Fig. 1B). In the data import step (`mdx2.import_data`), the diffraction images were re-compressed with three-dimensional chunks chosen to encompass each detector panel and 2° of rotation, or 20 images (see Supporting Information). Because subsequent processing steps distribute data among workers (CPU cores) according to chunk shape, the choice can affect performance. We have not systematically evaluated different chunk sizes, however the initial choice worked well in tests. The background images were also imported and re-binned every 10° and 20 pixels in each direction, reducing each stack of 360 images with 2463 × 2527 pixels to an array of 36 × 124 x 127 (`mdx2.bin_image_series`).

The next task in data processing was to accumulate the diffuse scattering on a three-dimensional grid. In order to mask out Bragg peaks, the coordinates of all pixels recording more than 20 photons (see Section 1.1.2) were fit to a three-dimensional Gaussian distribution in reciprocal space (mdx2.find_peaks). A peak mask was created at each Bragg peak location by thresholding the Gaussian distribution at 3 standard deviations (`mdx2.mask_peaks`). For integration, a grid was chosen that oversampled the reciprocal lattice by a factor of 3 in each direction (e.g., each voxel region spans Miller indices -1/6 to 1/6, 1/6 to 1/2, 1/2 to 5/6, etc). Because space group I 2_1_ 3 has the reflection condition *h* + *k* + *l* = 2*n*, only half of the voxels with all-integer Miller indices contain Bragg peaks, and thus for every Bragg peak there are 3^3^ · 2 − 1 = 17 voxels of diffuse scattering. Finally, integrated intensities were corrected for the experimentally-measured background scattering as well as polarization, air absorption, detector efficiency, and solid angle (`mdx2.correct`).

### 3.3 Scaling and analysis of systematic errors

In order to correct for systematic errors, a multi-crystal scaling model was refined (`mdx2.scale` with the flag`--mca2020`). As described in Section 2.1, this model accounts for changes in illuminated volume, excess background scattering, absorption of diffracted X-rays, and detector flat-field errors (Eq. 14 and Table 1). Default values were used for the grid dimensions, regularization parameters, outlier rejection thresholds, and stopping conditions (see Section 2.2 and the Supporting Information).

In general, the refined scaling model parameters offer insight into the types of systematic errors present during data collection. This is particularly true of the highly-redundant insulin dataset, because the model parameters are expected to be robustly determined. We examined each correction visually; representative sets are presented in Fig. 3. The overall scale factor for each frame varies in a similar manner to the corresponding Bragg scale factor refined in *DIALS* (Fig. S1 in the Supporting Information), which is significant because Bragg intensities are not included in the diffuse dataset refined in *mdx2*. Offset corrections varied greatly among the different wedges of data (Fig. 3A, Offset column). This variation is consistent with expectations; the large crystals were mounted with very little excess liquid on the surface (crystals in Fig. 2), and thus for most of the rotation range, the beam should pass only through the crystal without encountering the loop or excess solvent. The absorption corrections were relatively large for this dataset (Fig. 3A, Absorption column), with variation across the detector of as much as ±10%, which is expected given the exceptionally large crystals used.

**Figure 3:**
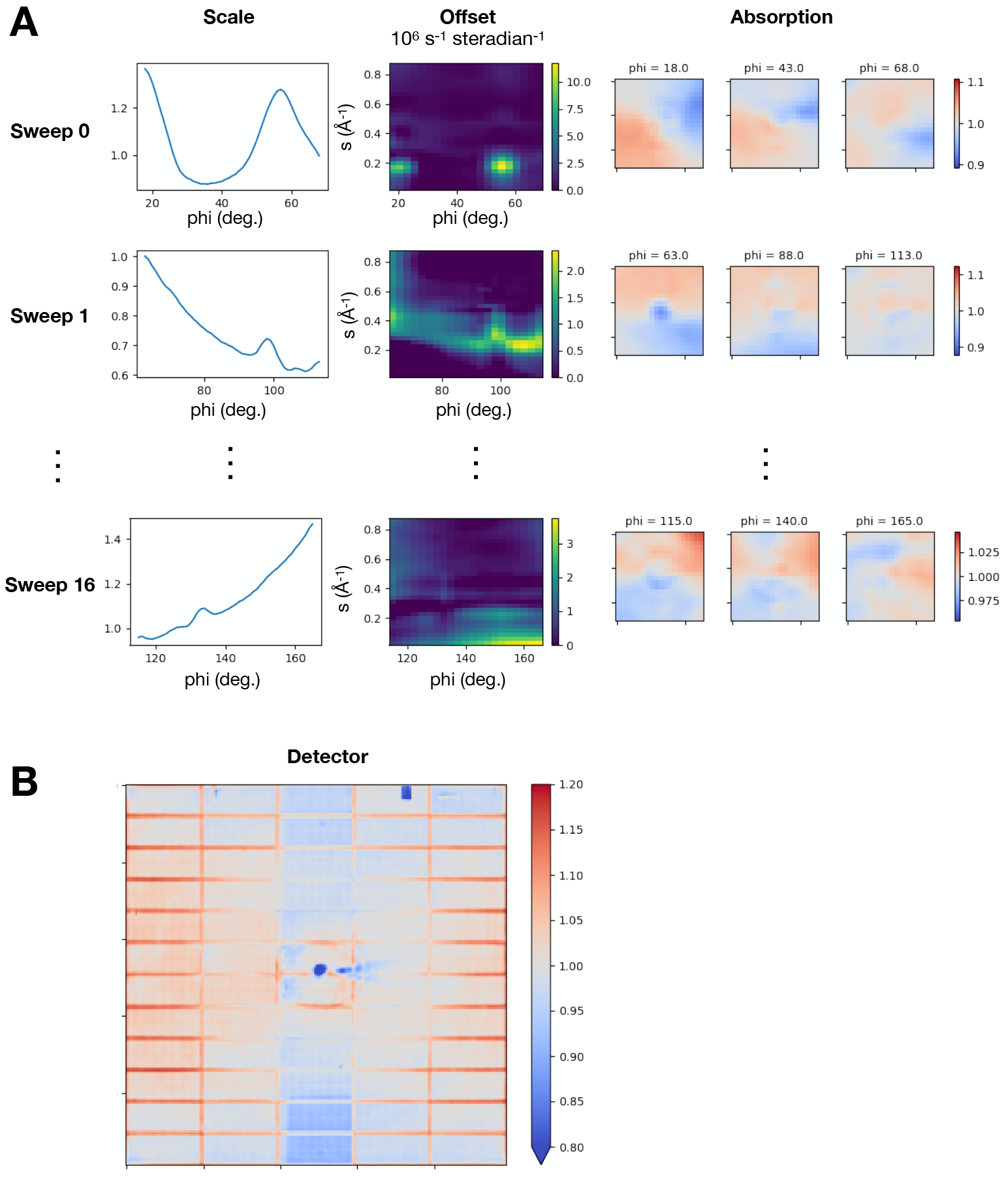
Diffuse scattering scaling model after refinement. The model was refined globally in *mdx2* to all 17 wedges in the multi-crystal insulin dataset. (**A**) Columns show each model parameter as a function of observation coordinates (rotation angle phi, detector position, and scattering vector magnitude, *s*). Each row corresponds to a 50-degree wedge (sweep) of data (only 3 are shown for clarity). Panels, from left to right, show the scale correction (*b* parameter), the offset correction (*c* parameter), and the absorption correction (*a* parameter) at the beginning, middle, and end of the rotation range. (**B**) The detector correction (*d* parameter) refined globally to all 17 wedges.

The detector flat-field correction was fit globally to all datasets. In contrast to the detector gain correction implemented previously in *mdx-lib*, where a single scale factor was applied for each detector chip (96 parameters), the flat-field correction in *mdx2* interpolates a grid covering the detector surface (200-by-200 = 40,000 parameters); the *mdx2* correction thus contains more potential detail about detector response than was available previously. Many of the detector errors were known in advance; for instance, one of the chips along the top row of the detector had malfunctioned and was reading lower values than the others. This chip is clearly visible in the corrections (blue rectangle in Fig. 3B, Detector). Another striking feature is the vertical stripe approximately of one panel width through the middle of the image. This feature results from absorption by the strip of Kapton film that suspends the beamstop. Finally, we note that data recorded at the edges of the detector panels are consistently lower intensity, and are boosted as much as ∼20% by the scaling correction (red outlines around each panel in Fig. 3B). After scaling, the redundant observations were merged according to the Laue symmetry (`mdx2.merge`).

### 3.4 Statistical analysis of intensities and data quality indicators

In all previously-collected datasets from lysozyme^14,20^, the voxels near the Bragg peaks contain more intense diffuse scattering (halos) compared with voxels further away. This phenomenon is expected to be a general feature of protein crystal diffuse scattering as it relates to the long-ranged coupling of subtle lattice motions^13,18,19,37^. Note that, in general, halo features can be broad and anisotropic depending on the correlation length of lattice motions. In lysozyme, halos tend to decay gradually according to the inverse-square of the distance from the Bragg peak^14,20^. To roughly quantify the halo scattering present in the insulin dataset, we split the merged data into two parts: the “halo” part consisted of only those voxels with integer Miller indices satisfying the reflection condition (*h*+*k*+*l* = 2*n*); and the rest, which we call “non-halo”. We do not expect the “non-halo” fraction for insulin to be completely free of scattering related to lattice disorder, however statistical analysis of intensities benefits from separately treating the voxels dominated by intense halo scattering. The “halo” voxels in this case correspond to a cubic region of reciprocal space with each side having Miller indices -1/6 to 1/6 with the central Bragg peak removed. The “halo” voxvels include fractional scattering vectors from (mean) radius of the peak mask at 0.102 to the corner of the voxel at 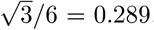. These vectors correspond to wave-like lattice displacements with wavelengths between 275–780 Å.

To better understand the components of the diffuse scattering signal, we computed the mean and standard deviation of the merged intensity within shells of constant resolution (Fig. 4A). The mean intensity rises to a peak at ∼3 Å resolution (Fig. 4A, top panel), a common feature of diffuse scattering from protein crystals originating from multiple sources: disordered solvent present in the pores of the crystal, uncorrelated atomic motion, and some short-range correlated motion^30,38^. The mean intensity for the halo fraction is greater than that of the non-halo fraction (Fig. 4A, top panel), consistent with the presence of diffuse scattering from lattice disorder. The intensity variations, containing the signal of greatest interest for studies of protein dynamics, can be quantified by the standard deviation of the intensity in each resolution shell. The variations are significantly larger for the halo fraction (Fig. 4A, bottom panel) compared with the non-halo part. The non-halo variations are small – less than 10% of the mean diffuse scattering in each resolution shell – underscoring the importance of careful measurement and scaling to determine this signal accurately.

**Figure 4:**
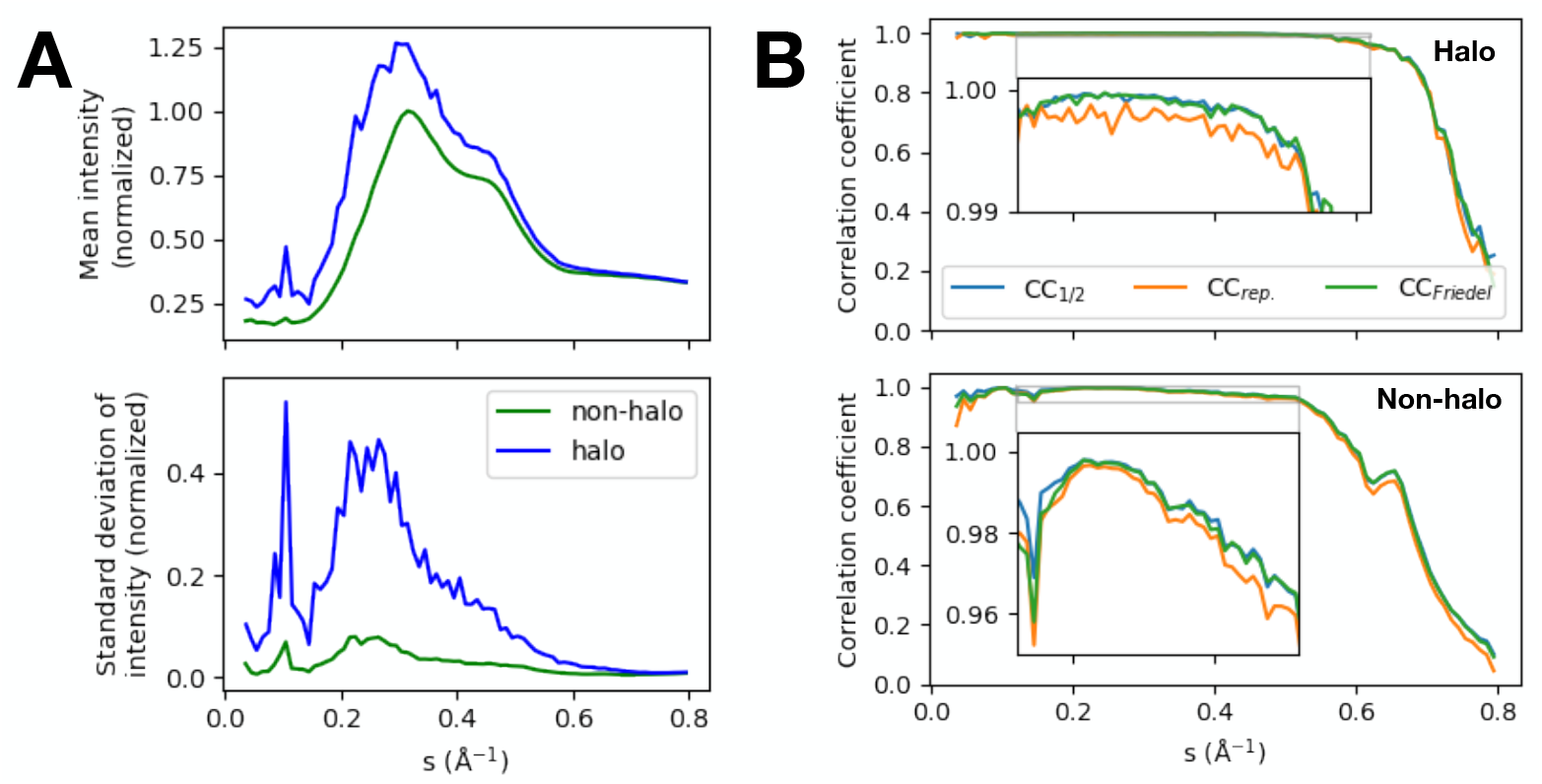
Diffuse intensity statistics and data quality indicators. (**A**) Merged intensities were split between halo- and non-halo voxels (see Main Text) and binned in shells of constant scattering vector *s*, where 1/*s* is the resolution. The mean intensity (top panel) rises to a peak at ∼3 Å resolution (∼0.3 Å^-1^). The intensities are normalized to 1 at the non-halo maximum. The halo-containing voxels have higher intensity on average than non-halo voxels. The intensity variations of interest are quantified by the standard deviation within each resolution shell (bottom panel). The halo-containing voxels have approximately 5-fold greater signal of non-halo voxels. Within each resolution shell, the non-halo variations are less than 10% of the mean scattering, indicating that this signal is much more subtle. (**B**) The correlation coefficient for random half-datasets (CC_1/2_), between crystals 1 and 2 scaled independently (CC_rep._), and between half-datasets split according to Friedel symmetry (CC_Friedel_) are compared for halo-containing voxels (top panel) and non-halo voxels (bottom panel). Correlation coefficients are close to 1 for much of the resolution range (insets), and decay at high resolution as the signal-to-noise ratio decreases. The correlations are higher overall for halo-containing voxels, as expected from the higher signal strength. There is no significant difference between random selection of observations (CC_1/2_) and grouping observations based on symmetry considerations (CC_Friedel_), which is expected for successful scaling. The CC_rep._ statistic closely follows CC_1/2_, showing that both the measurement and data processing procedures are reproducible.

The precision of diffraction data may be quantified by statistics reporting the agreement between equivalent observations. For diffuse scattering, we have previously quantified precision within each resolution shell using CC_1/2_, the Pearson correlation coefficient of intensity merged from random half-datasets^14,20^. To facilitate this calculation, a data splitting feature has been implemented in `mdx2.merge`, where various criteria may be used. To compute CC_1/2_, we split the data using a random shuffling algorithm that distributes equal numbers of equivalent observations between the two half-datasets (`--split randomHalf option`). The merged half-datasets are present as separate columns in the data table output by `mdx2.merge.` Because the scattering in halo voxels exceeds that of non-halo voxels, they will tend to dominate statistics such as CC_1/2_. Thus, the halo- and non-halo parts were analyzed separately. Correlation coefficients were close to 1 for both halo- and non-halo parts (Fig. 4B, blue lines in top and bottom panels respectively). In general, the halo-voxels have a slightly higher correlation than non-halo voxels at all resolutions, consistent with their greater signal-to-noise ratio. The signals are measured with very high precision in regions of high signal-to-noise: within all resolution shells up to 2 Å, CC_1/2_ exceeds 0.95 for non-halo and 0.99 for halo voxels. In both cases, CC_1/2_ decays at high resolution as the signal diminishes. However it remains significantly greater than zero up to 1.25 Å resolution (*s*<0.8 Å^-1^), even for the more subtle non-halo signal (Fig. 4B, bottom panel), suggesting that meaningful diffuse signal is present throughout reciprocal space.

A potential limitation of using the correlation between half-datasets to quantify precision is that such statistics may be inflated by certain systematic errors. A recent study compared various statistical measures of precision to address this possibility^22^. Here we follow a similar strategy. One alternative to the random split used in CC_1/2_ is to split according to Friedel symmetry (i.e., whether or not the symmetry operator mapping the observation to the asymmetric unit contains a center of inversion). Because equivalent observations differing by a center of inversion tend to be measured far apart in terms of their scan coordinates (e.g., detector position or sample rotation angle), the correlation coefficients from such a split will emphasize systematic differences in the data. To test this idea, we implemented such a split in `mdx2.merge`, which is activated by the `--split Friedel` option.

For the insulin dataset, we find that CC_Friedel_ and CC_1/2_ are indistinguishable across all resolution shells, both for the halo-containing voxels (Fig. 4B, top panel) and for the more subtle patterns in the non-halo voxels (Fig. 4B, bottom panel). The equivalence of CC_Friedel_ and CC_1/2_ would be expected if the scaling model corrects for all of the significant differences between equivalent observations, and thus it can be used to verify that a particular scaling model is sufficiently realistic.

A second, alternative splitting scheme is possible if multiple crystals are measured. In general, crystals have different shapes and are mounted in different orientations, and thus when comparing these independent datasets, there is less chance of spurious correlations arising from the chance alignment of a crystal symmetry with a particular geometric effect, such directional differences in absorption^22^. The correlation coefficient between independent datasets measures reproducibility, both of the measurement itself as well as the scaling procedure, and is called CC_rep._^22^.

We repeated the scaling and merging steps for each insulin crystal separately and computed CC_rep._, the correlation between independent measurements. We find that CC_rep._ is very close to CC_1/2_ at all resolution shells. When examined closely (Fig. 4B, insets), we find that CC_rep._ is always slightly below CC_1/2_, suggesting that CC_1/2_ overestimates reproducibility by a small margin. The reproducibility is still excellent; we attribute this result to the high redundancy of the insulin data and its highly symmetric Laue group, which ensures that the scaling model is tightly constrained by the data. Next, we investigated how data quality depends on the number of datasets that are merged. As expected when datasets are equivalent, CC_1/2_ improves at all resolutions as more datasets are added (Fig. S2 in the Supporting Information). The high multiplicity of the data was necessary in order to obtain good signal-to-noise for the non-halo data beyond ∼2 Å resolution (*s* > 0.5 Å^-1^). In summary, this comparison of intensity statistics suggest that meaningful diffuse signal is present at all resolutions and that systematic errors have been sufficiently corrected.

### 3.5 Visualizing weak features

To visualize the diffuse patterns, we symmetry-expanded and exported the merged data as a three-dimensional array (`mdx2.map`). A slice through this map containing only non-halo voxels (*l*=2/3) is shown in Fig. 5A. The pattern is dominated by the mostly-isotropic scattering. To better visualize the variational features, we subtracted the isotropic scattering. Here, we defined the isotropoic part as the mean non-halo scattering from each resolution shell interpolated at the reciprocal space coordinates of each voxel (a complete script is provided in the Supplementary Information). In this subtracted map, the variations are visible throughout (Fig. 5B).

**Figure 5:**
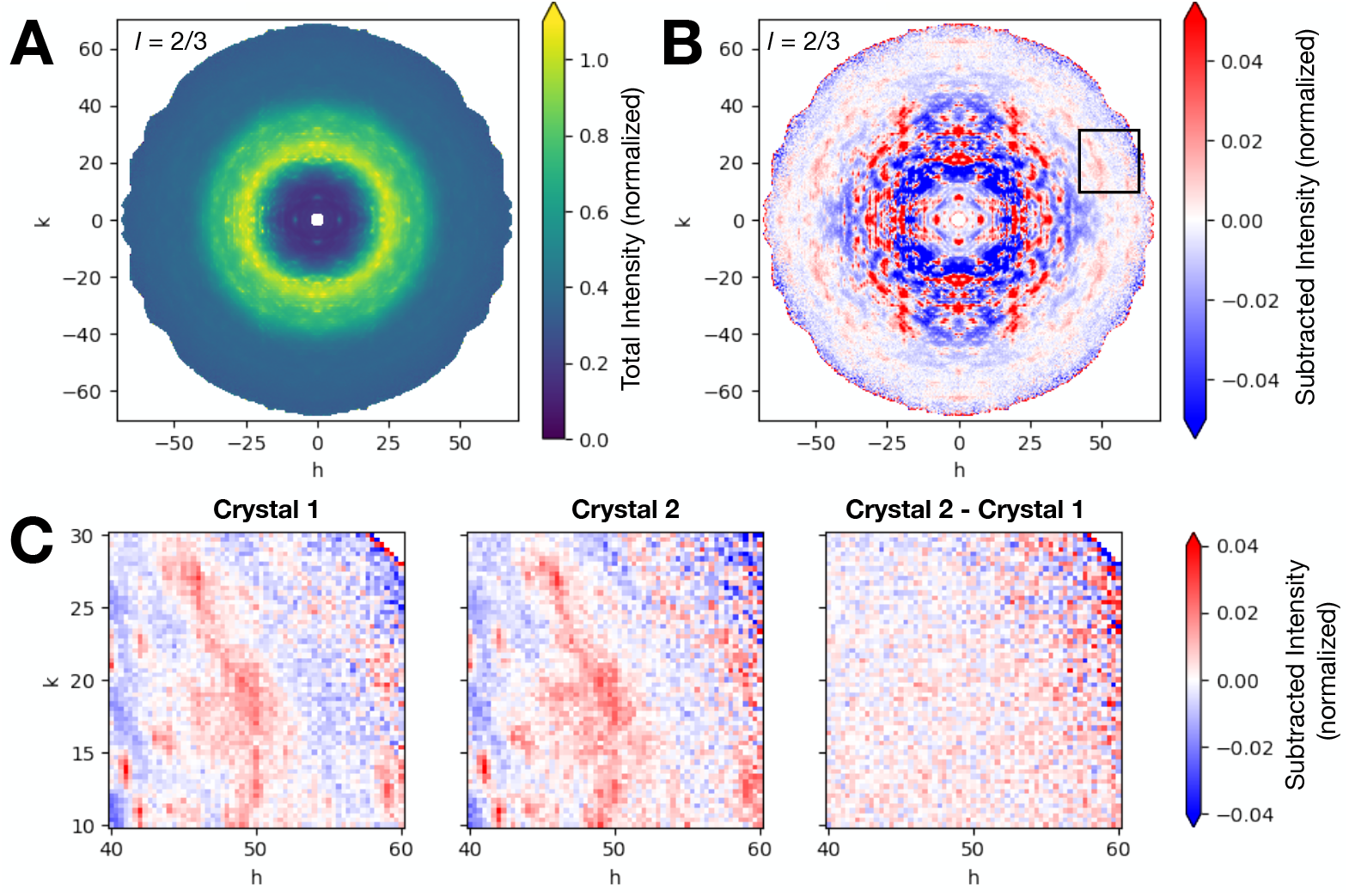
Visualization of weak diffuse scattering features. (**A**) A slice through the symmetry-expanded, three-dimensional map from cubic insulin in a non-halo plane (*l*=2/3). The intensity is normalized as in Fig. 4. (**B**) The same slice with the mean subtracted from each resolution shell to reveal subtle intensity variations. Features are visible at high resolution. (**C**) Comparison of the boxed region in panel B for two maps obtained by independently scaling and merging data from the two crystals. The pattern is reproduced in both maps, and the difference between the two appears random (right panel). The code used to generate this figure is provided in the Supplementary Information.

Our analysis of the data quality metrics, discussed above, suggest that the diffuse pattern at high resolution contains significant signal despite the measurement noise, and that it is uncorrupted by systematic errors. To verify that the patterns are indeed accurately determined, we compared maps that were scaled and merged separately from the two insulin crystals. As an illustrative example, we chose a small region containing a recognizable pattern (boxed region in Fig. 5B). Agreement between the reciprocal space maps of the two crystals is excellent, both in terms of the precise pattern and its overall magnitude (Fig. 5C, left and middle panels). Moreover, when one crystal map is subtracted from the other, the residual appears random (Fig. 5C, right panel). In general, such visual tests can be important to build confidence in a datasets’ accuracy, which is especially important when very subtle diffuse features such as these are used to support models of correlated motion.

## 4 Conclusions

We have described *mdx2*, a user-friendly software package for processing macromolecular diffuse scattering that incorporates state-of-the-art algorithms for data processing. A detailed scaling model, newly available in *mdx2* version 1.0, fully corrects for systematic errors in multi-crystal experiments. The complete processing of a multi-crystal dataset was demonstrated using the *mdx2* and *DIALS* command-line interfaces. This application is a clear example of how the close connection between *mdx2* and *DIALS* can be utilized to build complete workflows through simple scripting. At the same time, *mdx2* enables interactive data exploration through its commitment to the NeXus format for data interchange. *Mdx2* can also be imported as a Python toolkit for implementing custom algorithms.

The high-quality diffuse scattering maps generated for insulin highlight the potential benefits of high-redundancy data collection for diffuse scattering. When paired with the detailed scaling model in *mdx2*, redundancy can be leveraged to reduce the impact of systematic errors. Here we achieve ∼70-fold redundancy by collecting data from two large crystals in a high-symmetry space group. If crystals are low symmetry, or cannot be obtained with sufficient size, it may be necessary to collect data serially (i.e., from a large number of small crystals). Our results suggest that the high redundancy inherent to serial crystallography may be a useful feature for diffuse data processing, so long as background scattering can be minimized. We expect future versions of *mdx2* to include methods optimized for serial data.

The macromolecular diffuse scattering field is currently small, in part because substantial technical expertise has been required to process the data. However, based on the rapid progress enabled by modern detectors and computational methods, we can envision a future where diffuse scattering is part of the standard MX toolkit. Ultimately, diffuse scattering promises to enhance our understanding of protein dynamics and answer fundamental questions in biochemistry and biophysics. To achieve this goal, it is important to make diffuse scattering techniques accessible to researchers addressing important biological questions. *Mdx2* represents a step in this direction.

## Acknowledgments

We thank Neti Bhatt, Haoyue Wang, and Xiaokun Pei for testing the software and providing early feedback. The translation of sparse matrix interpolation code from MATLAB to Python followed the example performed by Darren Xu in the REGALS software package. We thank Kevin Dalton for getting us started with *dxtbx*, and the *DIALS* development team for answering questions when they arose during development. The software was run on shared computing resources available at CHESS, and we are greatful to the CLASSE computing group, especially Devin Bougie, for assistance with Python environments and for installing and maintaining a Jupyter Hub server that was used extensively for this project.

## Funding Information

Experiments were performed at the Center for High-Energy X-ray Sciences (CHEXS), which is supported by the National Science Foundation (BIO, ENG and MPS Directorates) under award DMR-1829070, and the Macromolecular Diffraction at CHESS (MacCHESS) facility, which is supported by award P30-GM124166 from the National Institute of General Medical Sciences, National Institutes of Health, and by New York State’s Empire State Development Corporation (NYSTAR). This work was supported by National Institutes of Health grant GM124847 (to N.A.).

## Data and Software Availability

The *mdx2* software package is available on GitHub (https://github.com/ando-lab/mdx2) and is free to use (GNU General Public License version 3). Version 1.0 described here has been archived on Zenodo (doi:10.5281/zenodo.10519719). The diffraction images are available for download from Zenodo (doi:10.5281/zenodo.10515006). Bash scripts for re-processing the data are included in the Supplementary Information. The figures can be reproduced by executing the Jupyter notebooks in the insulin-multi-crystal example included with the *mdx2* source code.

## Notes

https://github.com/ando-lab/mdx2

https://doi.org/10.5281/zenodo.10515006

### Competing Interest Statement

The authors have declared no competing interest.

