## Supplementary Information for "Scaling and merging macromolecular diffuse scattering with *mdx2*"

Steve P. Meisburger[a*] and Nozomi Ando[b*]

[a]Cornell High Energy Synchrotron Source, Cornell University, Ithaca, New York 14850, USA.

[b]Department of Chemistry and Chemical Biology, Cornell University, Ithaca, New York 14850, USA.

#### Contents

**1 Supplementary figures** **2**

**2 Scaling options in *mdx2*** **4**

**3 Insulin data processing scripts** **6**

**4 Insulin data analysis** **10**

### 1 Supplementary figures

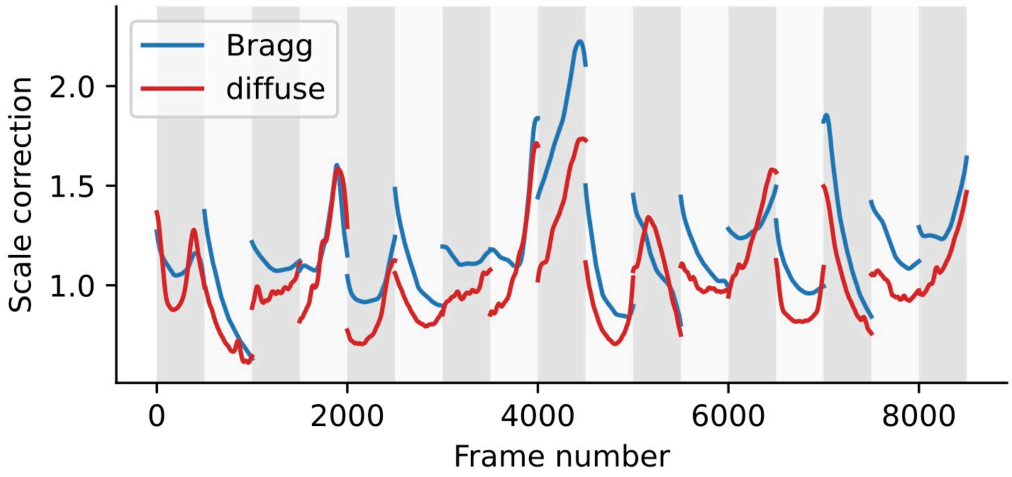


Figure S1: Per-image scale factors determined from Bragg data using *DIALS* (blue lines) and diffuse scattering data using *mdx2* (red lines). The data were re-plotted from the top axes in Fig. 2 and the Scale column in Fig. 3A for direct comparison (see Main Text). Each 50-degree wedge spans the width of the white/gray vertical bars. The scale factors show similar trends, but differ in detail.

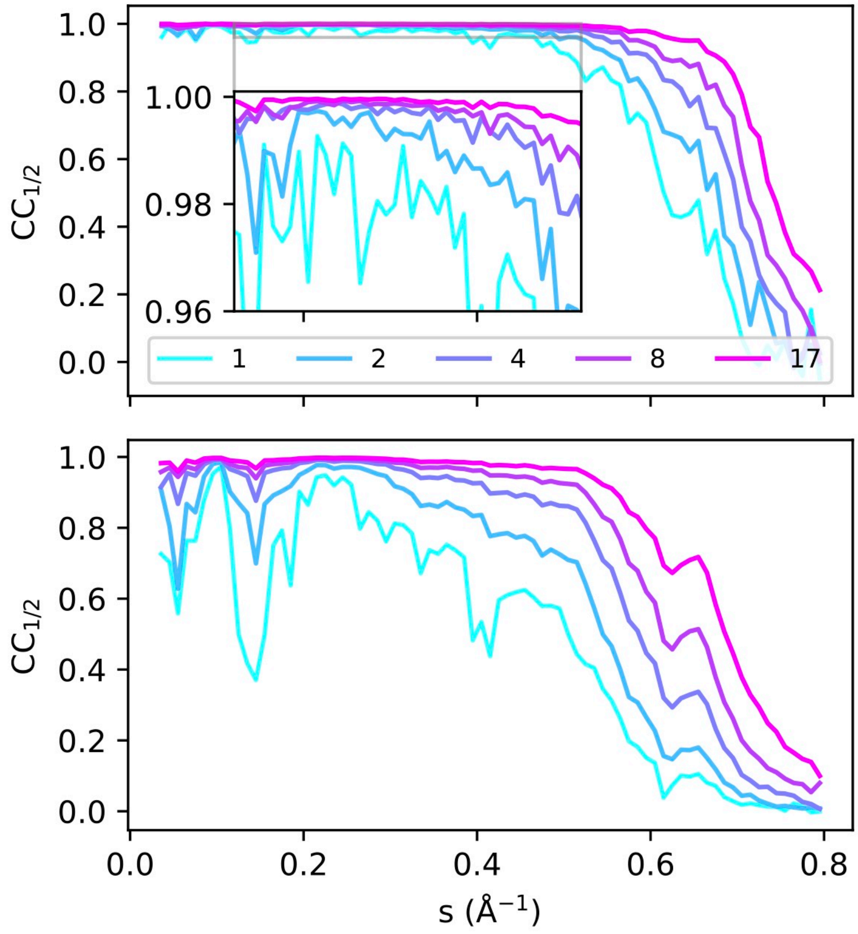


Figure S2: Improvement in diffuse data quality as datasets are added. The correlation coefficient of random half-datasets ($CC_{1/2}$) was calculated for partially merged data consisting of 1, 2, 4, 8, or 17 50-degree wedges. $CC_{1/2}$ was calculated separately for the halo-containing voxels (top panel) and the non-halo voxels (bottom panel), as explained in the Main Text.

#### 2 Scaling options in *mdx2*

A complete list of the command-line options for scaling in *mdx2* can be printed by running `mdx2.scale --help`. For reference, the output of this command (for mdx2 version 1.0) is reproduced below.

```
usage: mdx2.scale [-h] [--mca2020] [--outfile OUTFILE [OUTFILE ...]]
                  [--scaling.enable TF] [--scaling.alpha ALPHA]
                  [--scaling.dphi DEGREES] [--scaling.niter N]
                  [--scaling.x2tol TOL] [--scaling.outlier NSIGMA]
                  [--offset.enable TF] [--offset.alpha_x ALPHA]
                  [--offset.alpha_y ALPHA] [--offset.alpha_min ALPHA]
                  [--offset.min VAL] [--offset.dphi DEGREES] [--offset.ns N]
                  [--offset.niter N] [--offset.x2tol TOL]
                  [--offset.outlier NSIGMA] [--detector.enable TF]
                  [--detector.alpha ALPHA] [--detector.nx N] [--detector.ny N]
                  [--detector.niter N] [--detector.x2tol TOL]
                  [--detector.outlier NSIGMA] [--absorption.enable TF]
                  [--absorption.alpha_xy ALPHA] [--absorption.alpha_z ALPHA]
                  [--absorption.nx N] [--absorption.ny N]
                  [--absorption.dphi DEGREES] [--absorption.niter N]
                  [--absorption.x2tol TOL] [--absorption.outlier NSIGMA]
                  hkl [hkl ...]

Fit scaling model to unmerged corrected intensities

positional arguments:
  hkl                   NeXus file(s) with hkl_table

options:
  -h, --help            show this help message and exit
  --mca2020             shortcut for --scaling.enable True --offset.enable
                        True --detector.enable True --absorption.enable True
                        (default: False)
  --outfile OUTFILE [OUTFILE ...]
                        name of the output NeXus file(s). If not specified,
                        will attempt a sensible name such as scales.nxs for a
                        single input file (default: None)

Scaling parameters:
  --scaling.enable TF   include smooth scale factor vs. phi (default: True)
  --scaling.alpha ALPHA
                        amount to rescale the default smoothness
                        (regularization parameter) (default: 1.0)
  --scaling.dphi DEGREES
                        spacing of phi control points in degrees (default:
                        1.0)
  --scaling.niter N     maximum iterations in refinement (default: 10)
  --scaling.x2tol TOL   maximum change in x2 to stop refinement early
                        (default: 0.0001)
  --scaling.outlier NSIGMA
                        standard error cutoff for outlier rejection after
                        refinement (default: 10)

Offset parameters:
  --offset.enable TF    include smooth offset vs. resolution and phi (default:
                        False)
  --offset.alpha_x ALPHA
                        smoothness vs. s (resolution): multiplies the
                        regularization parameter (default: 1.0)
  --offset.alpha_y ALPHA
                        smoothness vs. phi -- multiplies the regularization
                        parameter (default: 1.0)
  --offset.alpha_min ALPHA
```

```
                        deviation from offset.min: multiplies the
                        regularization parameter (default: 0.001)
  --offset.min VAL      minimum value of offset (default: 0.0)
  --offset.dphi DEGREES
                        spacing of phi control points in degrees (default:
                        2.5)
  --offset.ns N         number of s (resolution) control points (default: 31)
  --offset.niter N      maximum iterations in refinement (default: 5)
  --offset.x2tol TOL    maximum change in x2 to stop refinement early
                        (default: 0.001)
  --offset.outlier NSIGMA
                        standard error cutoff for outlier rejection after
                        refinement (default: 5)

Detector parameters:
  --detector.enable TF  include smooth scale vs. detector xy position
                        (default: False)
  --detector.alpha ALPHA
                        smoothness vs. xy position: multiplies the
                        regularization parameter (default: 1.0)
  --detector.nx N       number of grid control points in the x direction
                        (default: 200)
  --detector.ny N       number of grid control points in the y direction
                        (default: 200)
  --detector.niter N    maximum iterations in refinement (default: 5)
  --detector.x2tol TOL  maximum change in x2 to stop refinement early
                        (default: 0.001)
  --detector.outlier NSIGMA
                        standard error cutoff for outlier rejection after
                        refinement (default: 5)

Absorption parameters:
  --absorption.enable TF
                        include smooth scale vs. detector xy position and phi
                        (default: False)
  --absorption.alpha_xy ALPHA
                        smoothness vs. xy position: multiplies the
                        regularization parameter (default: 10.0)
  --absorption.alpha_z ALPHA
                        smoothness vs. phi: multiplies the regularization
                        parameter (default: 1.0)
  --absorption.nx N     number of grid control points in the x direction
                        (default: 20)
  --absorption.ny N     number of grid control points in the y direction
                        (default: 20)
  --absorption.dphi DEGREES
                        spacing of phi control points in degrees (default:
                        5.0)
  --absorption.niter N  maximum iterations in refinement (default: 5)
  --absorption.x2tol TOL
                        maximum change in x2 to stop refinement early
                        (default: 0.0001)
  --absorption.outlier NSIGMA
                        standard error cutoff for outlier rejection after
                        refinement (default: 5)
```

### 3 Insulin data processing scripts

Data processing was orchestrated by chaining together *DIALS* and *mdx2* command-line programs using the Bourne Again Shell (BASH) scripting language. The scripts (lsts. 1, 2, 3, 4, 5, 6, 7, below) were executed sequentially on a 64-core linux server. If fewer cores are avilable, the parameter `--nproc` should be reduced. All scripts are run from the same base processing directory, and they produce the following output directory structure containing ~54 Gb of data:

```
. (project root)
├── dials
│   ├── 1_1
│   ├── 1_2
│   ├── 1_3
│   ├── 1_4
│   ├── 1_5
│   ├── 1_6
│   ├── 1_7
│   ├── 1_8
│   ├── 1_9
│   ├── 1_bkg
│   ├── 2_1
│   ├── 2_2
│   ├── 2_3
│   ├── 2_4
│   ├── 2_5
│   ├── 2_6
│   ├── 2_7
│   ├── 2_8
│   └── 2_bkg
└── mdx2
    ├── 1_bkg
    ├── 2_bkg
    ├── partial_merge
    ├── split_00
    ├── split_01
    ├── split_02
    ├── split_03
    ├── split_04
    ├── split_05
    ├── split_06
    ├── split_07
    ├── split_08
    ├── split_09
    ├── split_10
    ├── split_11
    ├── split_12
    ├── split_13
    ├── split_14
    ├── split_15
    └── split_16
```

**Listing 1** Bragg data processing

```
#!/bin/bash
set -e

DATADIR="/nfs/chess/scratch/user/spm82/data/insulin"
SUBS="1_1 1_2 1_3 1_4 1_5 1_6 1_7 1_8 1_9 2_1 2_2 2_3 2_4 2_5 2_6 2_7 2_8"
mkdir -p dials
cd dials
for sub in $SUBS; do
    mkdir -p $sub
    cd $sub
    dials.import "${DATADIR}/insulin_${sub}_*.cbf"
    dials.generate_mask imported.expt untrusted.circle=1264,1242,50
    dials.apply_mask imported.expt input.mask=pixels.mask output.experiments=imported.expt
    dials.find_spots imported.expt
    dials.index imported.expt strong.refl space_group=199
    dials.refine indexed.expt indexed.refl
    dials.integrate refined.expt refined.refl
    cd ..
done

dials.cosym **/integrated.{expt,refl} space_group=199
dials.scale symmetrized.{expt,refl} d_min=1.2
dials.split_experiments scaled.{expt,refl}
```

**Listing 2** Diffuse scattering integration

```
#!/bin/bash
set -e

mkdir -p mdx2
cd mdx2
for SUB in split_{00..16}; do
    mkdir -p $SUB
    cd $SUB
    EXPTFILE="../../dials/${SUB}.expt" # relative to subdir
    mdx2.import_geometry $EXPTFILE
    mdx2.import_data $EXPTFILE --chunks 20 211 493 --nproc 5 # single writer limits speed
    mdx2.find_peaks geometry.nxs data.nxs --count_threshold 20 --nproc 64
    mdx2.mask_peaks geometry.nxs data.nxs peaks.nxs --sigma_cutoff 3 --nproc 64
    mdx2.integrate geometry.nxs data.nxs --mask mask.nxs --subdivide 3 3 3 --nproc 64
    cd ..
done
```

**Listing 3** Background data processing (DIALS)

```
#!/bin/bash
set -e

DATADIR="/nfs/chess/scratch/user/spm82/data/insulin"

cd dials
for sub in {1..2}_bkg; do
    mkdir -p $sub
    cd $sub
    dials.import "${DATADIR}/insulin_${sub}_1_*.cbf"
    dials.generate_mask imported.expt untrusted.circle=1264,1242,50
    dials.apply_mask imported.expt input.mask=pixels.mask output.experiments=imported.expt
    cd ..
done
```

**Listing 4** Background data processing (mdx2)

```
#!/bin/bash
set -e

cd mdx2
for sub in {1..2}_bkg; do
    mkdir -p $sub
    cd $sub
    EXPTFILE="../../dials/${sub}/imported.expt"
    mdx2.import_data $EXPTFILE --chunks 10 211 493 --nproc 5
    mdx2.bin_image_series data.nxs 10 20 20 --valid_range 0 200 --outfile binned.nxs
    cd ..
done
```

**Listing 5** Diffuse intensity corrections

```
#!/bin/bash
set -e

for sub in split_{00..08}; do
    cd $sub
    mdx2.correct geometry.nxs integrated.nxs --background ../1_bkg/binned.nxs
    cd ..
done

for sub in split_{09..16}; do
    cd $sub
    mdx2.correct geometry.nxs integrated.nxs --background ../2_bkg/binned.nxs
    cd ..
done
```

**Listing 6** Scaling and merging

```
#!/bin/bash
set -e

cd mdx2

# scale all / merge all
mdx2.scale split_{00..16}/corrected.nxs --mca2020 --outfile split_{00..16}/scales_all.nxs
mdx2.merge split_{00..16}/corrected.nxs --scale split_{00..16}/scales_all.nxs \
  --outlier 5 --split randomHalf --outfile merged_all.nxs
mdx2.merge split_{00..16}/corrected.nxs --scale split_{00..16}/scales_all.nxs \
  --outlier 5 --split Friedel --geometry split_00/geometry.nxs --outfile merged_all_Friedel.nxs

# scale crystal 1 / merge crystal 1
mdx2.scale split_{00..08}/corrected.nxs --mca2020 --outfile split_{00..08}/scales_crystal1.nxs
mdx2.merge split_{00..08}/corrected.nxs --scale split_{00..08}/scales_crystal1.nxs \
  --outlier 5 --split randomHalf --outfile merged_crystal1.nxs

# scale crystal 2 / merge crystal 2
mdx2.scale split_{09..16}/corrected.nxs --mca2020 --outfile split_{09..16}/scales_crystal2.nxs
mdx2.merge split_{09..16}/corrected.nxs --scale split_{09..16}/scales_crystal2.nxs \
  --outlier 5 --split randomHalf --outfile merged_crystal2.nxs
```

**Listing 7** Partial merge (data for Fig. S2)

```
#!/bin/bash
set -e

cd mdx2
mkdir -p partial_merge

mdx2.merge split_00/corrected.nxs --scale split_00/scales_all.nxs \
  --outlier 5 --split randomHalf --outfile partial_merge/merged_1.nxs

mdx2.merge split_{00..01}/corrected.nxs --scale split_{00..01}/scales_all.nxs \
  --outlier 5 --split randomHalf --outfile partial_merge/merged_2.nxs

mdx2.merge split_{00..03}/corrected.nxs --scale split_{00..03}/scales_all.nxs \
  --outlier 5 --split randomHalf --outfile partial_merge/merged_4.nxs

mdx2.merge split_{00..07}/corrected.nxs --scale split_{00..07}/scales_all.nxs \
   --outlier 5 --split randomHalf --outfile partial_merge/merged_8.nxs

mdx2.merge split_{00..11}/corrected.nxs --scale split_{00..11}/scales_all.nxs \
  --outlier 5 --split randomHalf --outfile partial_merge/merged_12.nxs

mdx2.merge split_{00..16}/corrected.nxs --scale split_{00..16}/scales_all.nxs \
  --outlier 5 --split randomHalf --outfile partial_merge/merged_17.nxs

cd ..
```

### 4 Insulin data analysis

Statistical analysis of scattering and visualization tasks were performed in Python using the *mdx2* package and standard tools such as *pandas* and *numpy*. All pre-processing and plotting steps for Figures 2–5 in the Main Text, and Figures S1, and S2 are encapsulated in Jupyter notebooks that are distributed with *mdx2*: `https://github.com/ando-lab/mdx2/tree/main/examples/insulin-multi-crystal`. As an illustrative example, we include here the scripts used to calculate and subtract the isotropic component of scattering (lst. 8), generate slices through the three-dimensional map (lst. 9), and create the panels in Figure 5 in the Main Text (lst. 10).

**Listing 8** Calculate the isotropic component and subtract

```
from mdx2.utils import loadobj, saveobj
import pandas as pd
import numpy as np

Crystal  = loadobj('mdx2/split_00/geometry.nxs','crystal')
Symmetry = loadobj('mdx2/split_00/geometry.nxs','symmetry')

def hkl2s(h,k,l):
    """Compute the magnitude of s from Miller indices."""
    UB = Crystal.ub_matrix
    s = UB @ np.stack((h,k,l))
    return np.sqrt(np.sum(s*s,axis=0))

def isInteger(h):
    return (h.round()-h).abs() < 0.00001

def isReflection(h,k,l):
    return isInteger(h) & isInteger(k) & isInteger(l) & Symmetry.is_reflection(h,k,l)

def calc_stats(fn):
    tab = loadobj(fn,'hkl_table')
    tab.s = hkl2s(tab.h,tab.k,tab.l)
    df = tab.to_frame()
    df = df[~isReflection(df['h'],df['k'],df['l'])]
    df = df.set_index(['h','k','l']).sort_index()
    s_bins = pd.cut(df['s'],np.linspace(0,.8,81))
    df_isoavg = df.groupby(s_bins).agg({
        's':'mean',
        'intensity':['mean','std'],
        'intensity_error':'mean'})
    dfh = tab.to_frame()
    dfh = dfh[isReflection(dfh['h'],dfh['k'],dfh['l'])]
    dfh = dfh.set_index(['h','k','l']).sort_index()
    s_bins = pd.cut(dfh['s'],np.linspace(0,.8,81))
    dfh_isoavg = dfh.groupby(s_bins).agg({
        's':'mean',
        'intensity':['mean','std'],
        'intensity_error':'mean'})
    df_out = pd.DataFrame({
        's':df_isoavg['s']['mean'],
        'non-halo':df_isoavg['intensity']['mean'],
        'halo':dfh_isoavg['intensity']['mean']})
    df_out = df_out.set_index('s')
    return df_out

for dset in ['all','crystal1','crystal2']:
    df_stats = calc_stats(f'mdx2/merged_{dset}.nxs')
    nh = df_stats['non-halo'].dropna()
    x = nh.keys().values
    y = nh.values
    Imax = np.max(y)
    tab = loadobj(f'mdx2/merged_{dset}.nxs','hkl_table')
    tab.isoavg = np.interp(hkl2s(tab.h,tab.k,tab.l),x,y/Imax)
    tab.intensity/=Imax
    tab.intensity_error/=Imax
    tab.intensity -= tab.isoavg
    saveobj(tab,f'merged_{dset}_sub.nxs','hkl_table')
```

**Listing 9** Create three-dimensional arrays for visualization

```
mdx2.map mdx2/split_00/geometry.nxs merged_all_sub.nxs --limits -70 70 -70 70 0 1 \
  --signal isoavg --outfile slice_all_isoavg.nxs
mdx2.map mdx2/split_00/geometry.nxs merged_all_sub.nxs --limits -70 70 -70 70 0 1 \
  --signal intensity --outfile slice_all.nxs
mdx2.map mdx2/split_00/geometry.nxs merged_crystal1_sub.nxs --limits -70 70 -70 70 0 1 \
  --signal intensity --outfile slice_crystal1.nxs
mdx2.map mdx2/split_00/geometry.nxs merged_crystal2_sub.nxs --limits -70 70 -70 70 0 1 \
  --signal intensity --outfile slice_crystal2.nxs
```

**Listing 10** Plots for Figure 5 in the Main Text

```
import matplotlib.pyplot as plt
import xarray as xr
from nexusformat.nexus import nxload

imav = nxload('slice_all_isoavg.nxs')['/entry/isoavg']
imsub = nxload('slice_all.nxs')['/entry/intensity']
imsub1 = nxload('slice_crystal1.nxs')['/entry/intensity']
imsub2 = nxload('slice_crystal2.nxs')['/entry/intensity']

im = imsub + imav

def map2xr(m):
    return xr.DataArray(
        data=m.signal.nxvalue,
        dims=m.axes,
        coords=dict(h=m.h.nxvalue, k=m.k.nxvalue, l=m.l.nxvalue)
    )

# Figure 5, panels A and B
fig,(ax1,ax2) = plt.subplots(1,2,figsize=(10,4))
map2xr(im).isel(l=2).plot(x='h',y='k',ax=ax1,vmin=0,vmax=1.1,cmap='viridis')
map2xr(imsub).isel(l=2).plot(x='h',y='k',ax=ax2,vmin=-.05,vmax=.05,cmap='bwr')
ax1.set_aspect('equal')
ax2.set_aspect('equal')
plt.savefig('fig5ab.png',transparent=True)

# Figure 5, panel C
a = map2xr(imsub1).isel(l=2)
b = map2xr(imsub2).isel(l=2)
d = xr.concat((a,b,b-a),pd.Index(['1','2','2-1'],name="Crystal"))
fh = d[:,330:391,240:301].plot(x='h',y='k',col='Crystal',vmin=-.04,vmax=.04,cmap='bwr')
[ax.set_aspect('equal') for ax in fh.axs.flatten()]
plt.savefig('fig5c.png',transparent=True)
```
